# Bacteriophages inject DNA into giant unilamellar vesicles mimicking Gram-negative lipopolysaccharide outer membranes

**DOI:** 10.1101/2024.08.19.608551

**Authors:** Mareike S. Stephan, Nina K. Broeker, Tom Robinson, Rumiana Dimova, Stefanie Barbirz

**Affiliations:** Max Planck Institute of Colloids and Interfaces, Potsdam, Germany; Department of Medicine, Science Faculty, HMU Health and Medical University, Potsdam, Germany; Department of Medicine, Science Faculty, MSB Medical School Berlin, Germany; Institute for Bioengineering, School of Engineering, University of Edinburgh, United Kingdom

**Keywords:** giant unilamellar vesicles, model membrane, bacteriophage, lipopolysaccharide, DNA transfer

## Abstract

Giant unilamellar vesicles (GUVs) are a versatile platform to study cell membrane functions. We have constructed phospholipid GUVs presenting lipopolysaccharide on their external surface (LPS-GUVs) to mimic the outer membrane (OM) of Gram-negative bacteria. GUVs allow for adjusting a defined OM composition, unlike the dynamic changes of LPS structures typically observed in vivo. The OM is a major control point for the viral genome transfer from bacteriophages into bacterial hosts. We found that siphovirus 9NA specifically binds to the surface of GUVs when presenting its *Salmonella* Typhimurium LPS phage receptor. Using LPS-GUVs filled with DNA-sensitive dyes we show that after surface fixation, the bacteriophage particle opens and injects its DNA into the GUV lumen. No OM proteins were included in the LPS-GUV membrane, emphasizing that the presence of the LPS membrane glycolipid assembly alone is sufficient to trigger the start of bacteriophage genome transfer. LPS-GUVs thus open a sustainable route to systematic studies of viral infection mechanisms at the host envelope and provide a cell-free platform to study surfaces of pathogenic bacteria. This is an important prerequisite for developing effective antimicrobial therapies based on bacteriophages that target Gram-negative pathogens.

---

Bacterial pathogens have emerged as a major global health threat with increasing treatment resistances, especially to last-resort antibiotics that target Gram-negative bacteria.^1-2^ Bacteri-ophages, i.e. viruses that prey on bacteria, are considered highly important to be included in future antimicrobial strategies.^3^ Their effectiveness requires fundamental research on the mechanisms of bacteriophage action. Transit through the bacterial envelope is a key step in the lytic bacteriophage life cycle, eventually leading to viral genome transfer, intracellular replication and killing of the host bacterium prior to release of the progney.^4^ Here, we show that giant unilamellar vesicles (GUVs) are a versatile, cell-free platform to study bacteriophage genome transfer through outer membranes (OMs) of Gram-negative bacteria. We used GUVs that mimic the asymmetric Gram-negative OM^5^ and expose lipopolysaccharide (LPS) as bacteriophage membrane receptors on their outer surface. This setup allowed us to monitor phage DNA delivery into the GUV lumen with fluorescence microscopy (Fig. 1).

**FIGURE 1:**
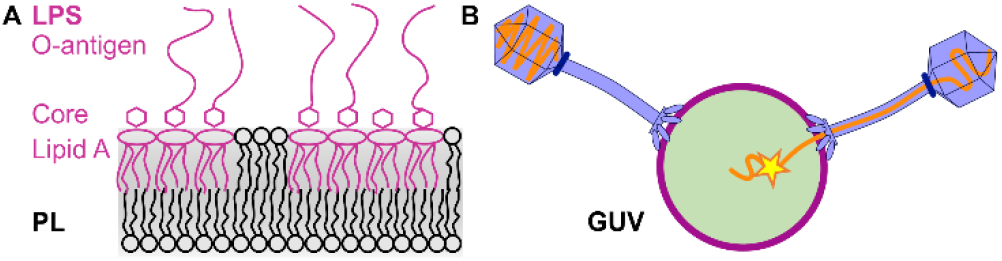
Schematic experimental set up. A) GUV phospholipid (PL) membrane exposing LPS (magenta) on the outer leaflet mimicking the Gram-negative bacterial envelope. B) Bacteriophages bind to the LPS receptor on GUVs and inject their DNA-genomes (orange), here monitored with fluorescent dye YO-PRO™-1 in the GUV lumen that detects transferred DNA (yellow).

GUVs have proven to be robust biomimetic systems for studying eukaryotic cell membrane functions and serve as intracellular delivery and protocell models.^6-8^ More recently, protocols for the formation of GUV models of the Gram-negative OM have been described.^5, 9-11^ Production of these GUVs is challenging because they must mimic a heterobilayer composed of an intracellular phospholipid and an extracellular LPS leaflet (Fig. 1A).^12-13^ LPS, also known as endotoxin due to its strong immunogenic properties, is a complex glycolipid and serves as an important bacteriophage adsorption and infection mediator (Fig. 1B).^4, 14-15^ *In vitro*, isolated LPS molecules, typically form multilamellar, heterogeneous aggregates^14^ when they lack the ordered membrane environment of the living cell. Therefore incorporating LPS into biomimetic systems of the Gram-negative outer bacterial envelope is more challenging compared to using standard membrane phospholipids found in many cells.^16-17^

Tailed phages are the most abundant phage type isolated from bacterial ecosystems and antimicrobial therapy approaches typically employ bacteriophages that have tails and double stranded DNA.^3^ Tailed bacteriophages remain extracellular and only transfer their DNA through the cellular envelope.^18^ After adsorption to the bacterial envelope, the phage tail receives a specific signal from the cell surface, triggering conformational rearrangements in the tail machinery that opens the phage particle for subsequent genome ejection.^20^ For the initiation of infection, many bacteriophages interact with LPS that is often decorated with O-antigen polysaccharides to which phages can specifically bind. However, studying the role of LPS in mechanisms of phage DNA transfer into a bacterial cell is difficult *in vivo* because the OM composition dynamically changes throughout the bacterial life cycle, particularly during phage infection.^21^

We used LPS-GUVs as mimics of the Gram-negative OM to study bacteriophage genome transfer in a defined *in vitro* prokaryotic membrane system. To study bacteriophage interactions, LPS-containing GUVs must be clearly asymmetric, i.e. the double membrane must have phospholipids facing the GUV lumen whereas the phage LPS receptors are exposed on the GUV surface (Fig. 1).^5^ Moreover, the external GUV solution must be free of unbound LPS aggregates which compete with LPS-GUVs for phage binding.^22^ We chose *Salmonella* siphovirus 9NA as a model phage.^23^ 9NA is O-antigen specific, i.e. a smooth LPS containing the O-polysaccharide is a prerequisite to start DNA release from the phage and initiate *Salmonella* infection. We prepared asymmetric GUVs exposing LPS of *Salmonella* Typhimurium (*S*. Typhimurium) using an inverted emulsion technique as described previously (see SI).^5^ We employed microfluidics to allow both washing/purification of GUVs and controllable delivery of bacteriophages (see SI).

*In vitro*, protein-free LPS aggregates are sufficient to trigger full genome ejection from 9NA in a short time, even when not organized in a bacterial OM heterobilayer (Fig. S1).^23^ We therefore applied microfluidics for washing the GUVs to remove free and unbound LPS (see SI). This ensured that phages could predominantly interact with LPS on the GUV surface, and drastically reduced DNA ejection upon contact with residual LPS in solution. Phages were stained with the DNA-intercalating dye SYBR-gold, LPS was labelled with TexasRed or Alexa647.^10^

Bacteriophage 9NA specifically bound to the GUV outer surface via its LPS O-antigen receptor (Fig. 2 A-C). In contrast, 9NA phages did not localize to the GUV membrane when their O-antigen receptor was removed by incubating the GUVs with P22 tailspike, a phage enzyme that specifically cleaves off the *S*. Typhimurium O-antigen from LPS^19^ (Fig. 2 D-F). Bacteriophage 9NA did not bind to POPC GUVs that did not contain LPS (Fig. S2). The number of phages localized onto the GUV membrane scaled with the LPS concentration on the vesicle surface (Figs. 3 and S3). After binding to LPS via its tailspikes^23^, we observed a stable 9NA bacteriophage-binding signal at the membrane for the entire observation time (1 h). We hence assume that the 9NA tail machinery had reached the membrane surface and remained irreversibly bound. Bacteriophage 9NA can specifically distinguish *Salmonella* hosts from other nonhost strains, i.e. *E. coli*.^24^ Accordingly, 9NA did not bind to GUVs when prepared with LPS from an unrelated, O-antigen containing *E. coli* strain (Fig. S2). Moreover, 9NA did not bind to GUVs with inverted membranes - where phospholipids are on the outer leaflet and LPS is on the inside - and was unable to access the luminal LPS O-antigen receptors (Fig. S2). This underscores that our preparation method enabled precise tuning of the membrane asymmetry of LPS-GUVs.

**FIGURE 2.**
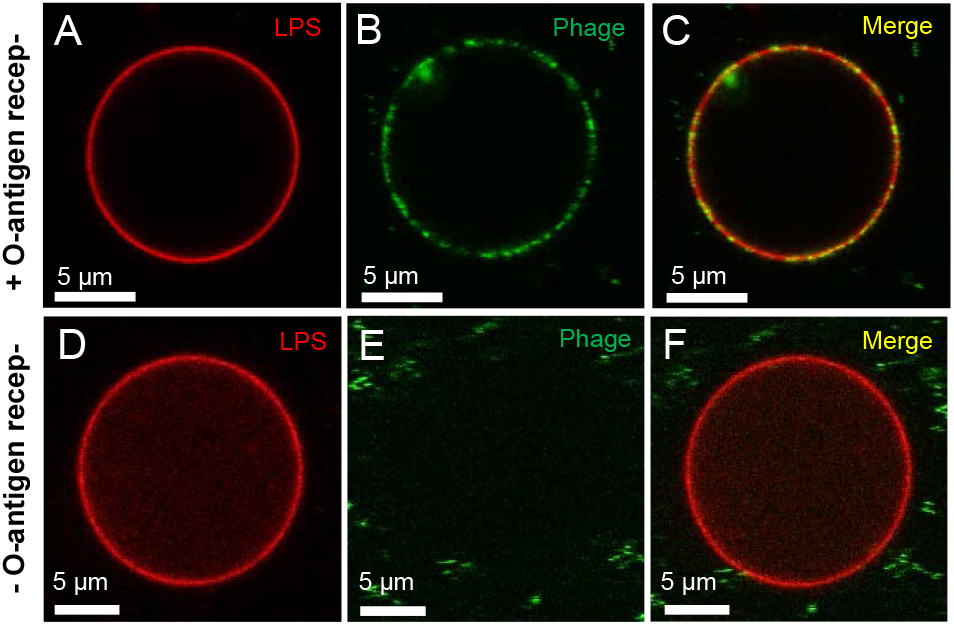
*Salmonella* siphovirus 9NA binds specifically to LPS-GUVs presenting O-antigen receptor on the outer surface. (A,D) TexasRed labelled *S*. Typhimurium LPS, (B,E) SYBR™Gold labelled phages (green), (C,F) overlaid signals. (D-F) Phage O-antigen receptor was enzymatically cut from surface LPS by incubating the GUVs with P22 TSP endorhamnosidase^19^ (25 μg ml^-1^, 25°C, overnight).

**FIGURE 3:**
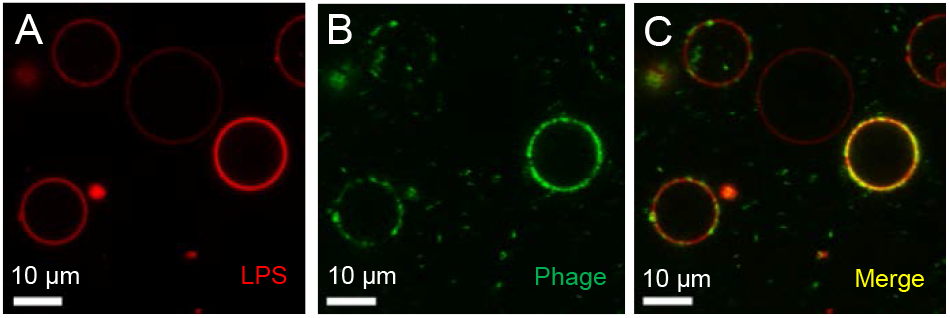
The amount of bound phages on the GUV surface increases with LPS concentration. LPS (A), phage (B) and merged signals (C) (*cf*. Fig. 2). For fluorescence intensity quantification see Fig. S3.

Over time after phage addition, the number of phage particles associated with the GUV membrane increased, and local variations in fluorescence intensity indicated that particles had ejected their DNA. However, due to the immediate dilution of SYBR gold upon ejection from the phage particle, we were unable to clearly resolve individual ejection events or determine the precise localization of the ejected DNA. Therefore, we prepared GUVs filled with TOTO™-3, a membrane-impermeable cationic DNA dye that also slightly binds and stains the membrane (Fig. 4). Immediately upon addition, we observed the binding of SYBR gold labelled 9NA phages to the GUV membrane, accompanied by single bright spots in the TOTO™-3 channel. This demonstrated the co-localization of membrane-associated viral particles and their ejected DNA.^25^

**FIGURE 4:**
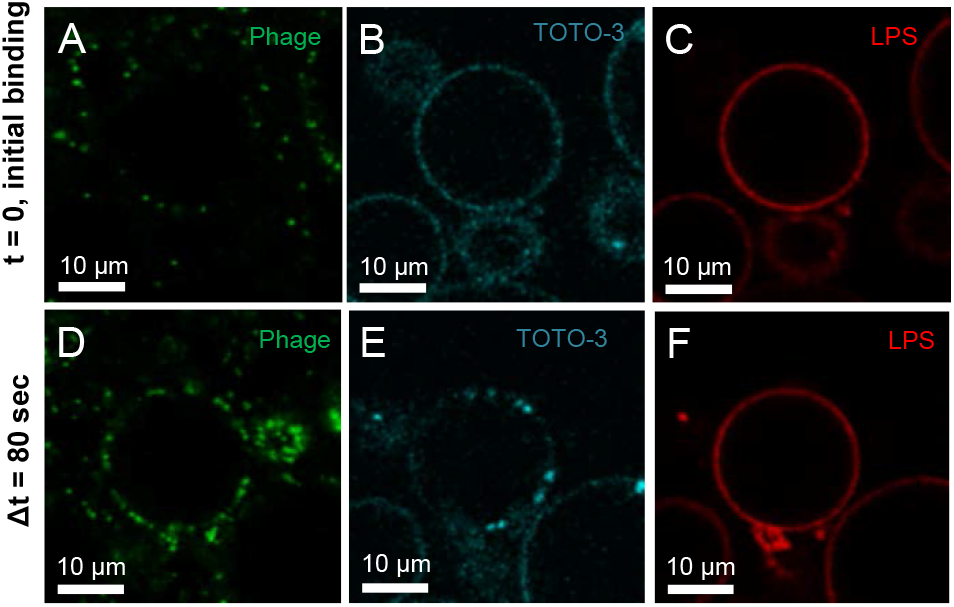
DNA signal observed at the surface of the same LPS-GUV upon bacteriophage binding. (A,B) SYBR-Gold labelled 9NA phages, (B,E) TOTO™-3 in solution inside LPS-GUVs. (C,F) LPS labelled with TexasRed. Images at time point of first observed 9NA phage binding events (A-C) and after Δt=80 s (D-F).

To unambiguously confirm that phage 9NA DNA was directed into the GUV lumen and did not remain in the extravesicular space, we used GUVs filled with the cell-impermeant cationic cyanine dye YO-PRO™-1 (Fig. 5). Whereas TOTO is a dimer, the monomeric YO-PRO has fewer charges and does not bind to the GUV membrane. Within 20 min of observation, an increasing number of bright spots appeared at the inner membrane leaflet, indicating DNA release from the phages into the GUV lumen (Fig. 5A and Movie S1). Variability in the number of ejecting phages among GUVs was observed, likely due to the limitation of rapid mixing in our set up, hampering clear time-resolved observations of ejection initiation. Instead, phages had to diffuse toward GUVs settled at the end of the microfluidic wells (see SI). Ejection events frequently occurred at various positions on the GUV surface (Fig. 5B). The DNA remained attached to the inner side of the GUV membrane; we observed all DNA as bright spots tethered to the inner leaflet, extending to approximately 3 μm into the GUV lumen. This irreversible attachment aligns with findings from cryo electron tomography, where phages were observed fixed to the OM via protein channels built for genome delivery.^20^

**FIGURE 5:**
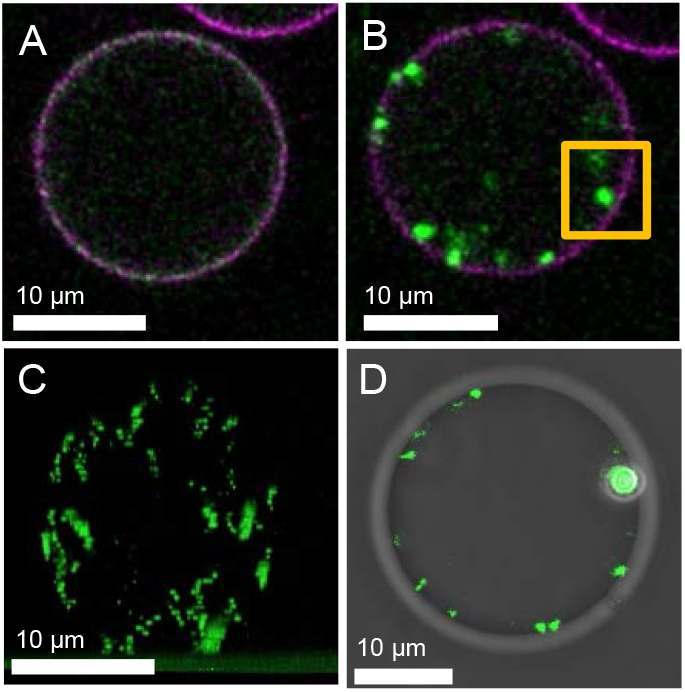
9NA bacteriophage injects its DNA into the LPS-GUV lumen. GUVs contained *S*. Typhimurium LPS (Alexa647-labelled, magenta) on the outer surface and 10 μM of DNA intercalating dye YO-PRO™-1 (green) in the lumen. (A) Snapshots of YO-PRO™-1 fluorescence, inset in (B) shows DNA signal located at the inner GUV membrane. (C) Z-stack of DNA signal after 15 min. (D) Phase contrast of an LPS-GUV after 25 min of incubation with 9NA phage. All measurements in presence of 0.5 mg ml^-1^ DNAse at room temperature.

*In vitro*, the release of densely packed DNA from phage capsids is driven solely by external osmotic pressure^26^. At osmotic equilibrium, typically 20 % of the DNA remain inside the phage heads^27^, whereas *in vivo* intracellular host protein interactions result in pulling the complete phage genome into the cell^28^. Therefore, in the absence of these factors, we observe only partially ejected phages tethered to the GUV membrane. After DNA transfer, any membrane poration was sealed and the membrane itself remained fully intact, as evidenced by the maintained phase contrast of the vesicles (Fig. 5C).

GUVs have been used so far as models to study the initial steps of virus endocytosis into eukaryotic hosts.^29-30^ In contrast, bacteriophages are not endocytosed but employ ejection proteins to build membrane-crossing channels and maintain envelope integrity during genome transfer into a bacterial host.^18, 31-32^ We could show that LPS-GUVs are an *in vitro*-mimic for this phage-based DNA transfer into bacteria. The LPS-containing GUV membrane was sufficient to direct phage DNA into the lumen while the GUV remained intact (Fig. 4C). In particular, *Salmonella* phage 9NA only needed an O-antigen receptor on the LPS, and no protein. This is in contrast to other siphoviruses, *i*.*e*. phages with long, non-contractile tails like lambda or T5 that employ OM proteins as receptors.^33-34^ T5 initiates formation of a DNA-transfer channel by docking next to its OM receptor FhuA as shown with nanodiscs and cryo electron microscopy.^33^ Furthermore, DNA transfer is possible into GUVs that had the FhuA receptor reconstituted in the membrane.^35^ However, neither the GUVs nor the nanodiscs contained LPS in these studies, despite LPS being responsible for the unique properties of the Gram-negative OM heterobilayer. In particular, it has been shown that especially long O-antigen chains on LPS contribute to local membrane rigidity which has been proposed as essential feature for infection start in O-antigen specific phages.^13, 22, 36^ We calculated diffusion coefficients from individual 9NA particle tracks when bound to LPS-GUVs to around 1.5 μm^2^ s^-1^. This value remained constant for the given range of varying LPS content in individual GUVs (Fig. S4). In contrast, phage T5, tethered to GUVs via its protein receptor FhuA and of similar size, diffused notably more rapidly on the LPS-free GUV at similar temperatures.^35^ Whether the irreversibly membrane-bound 9NA phage thus moved on the surface as part of a larger LPS aggregate or raft needs further investigation.

LPS is the essential OM glycolipid exposed on every Gramnegative cell, making LPS-GUVs highly versatile in vitro platforms to study bacteriophage interactions. We introduce a new model system that enables biophysical studies of virus-host membrane interactions without the need to handle pathogenic bacteria. It can be a starting point for a variety of applications, for example for DNA sorting from viral mixtures or for vesicle-based delivery of viral DNA for in-vesicle transcription-translation.^37^ Most importantly, successful therapies based on bacteriophages require that we understand the bacteriophage infection mechanism and why phage mixtures are superior to single phage applications.^3^ LPS-GUVs open new sustainable routes to fundamental mechanistic studies on infection synergies in the presence of different phage receptors using LPS preparations *in vitro*, and enable imaging phage DNA transfer via OMs of Gram-negative pathogens in a controlled in vitro set up.

## ASSOCIATED CONTENT

Supporting Information available: Additional experimental details, materials, and methods (PDF). Movie of 9NA bacteriophage DNA ejection into GUVs (MP4).

## AUTHOR INFORMATION

### Author Contributions

RD, TR and SB proposed and supervised the project. RD, TR, SB, NB and MS designed the experiments. MS performed the experiments and analyzed the data. RD, TR, SB, NB and MS wrote the manuscript.

## ACKNOWLEDGMENT

MS acknowledges funding from the International Max Planck Research School on Multiscale BioSystems. TR acknowledges funding from the MaxSynBio consortium, which is jointly funded by the Federal Ministry of Education and Research of Germany and the Max Planck Society.

## ABBREVIATIONS

GUV: giant unilamellar vesicles
LPS: lipopolysaccharide
OM: outer membrane
PL: phospholipid.

## Notes

### Competing Interest Statement

The authors have declared no competing interest.

