## Supplementary data for "Bacteriophages inject DNA into giant unilamellar vesicles mimicking Gram-negative lipopolysaccharide outer membranes"

For

### Supplementary Figures

#### Figure S1. LPS triggers DNA release from *Salmonella* bacteriophage 9NA

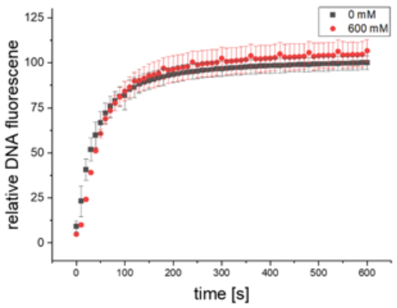

*Salmonella* bacteriophage 9NA DNA release is triggered by LPS in presence or absence of sucrose.[1] DNA ejection was followed by the detection of the fluorescence signal increase upon the incubation of $4*10^9$ pfu $ml^{-1}$ 9NA phages with 10 µg $ml^{-1}$ LPS solution at 37 °C in the presence of 1 µM of the fluorescent DNA-binding dye YO-PRO™-1. Error bars represent standard deviation of three independent experiments. Red curve in the presence of 600 mM sucrose.

#### Figure S2. *Salmonella* bacteriophage 9NA interactions with membrane-inverted GUVs, *E. coli* LPS GUVs and POPC GUVs

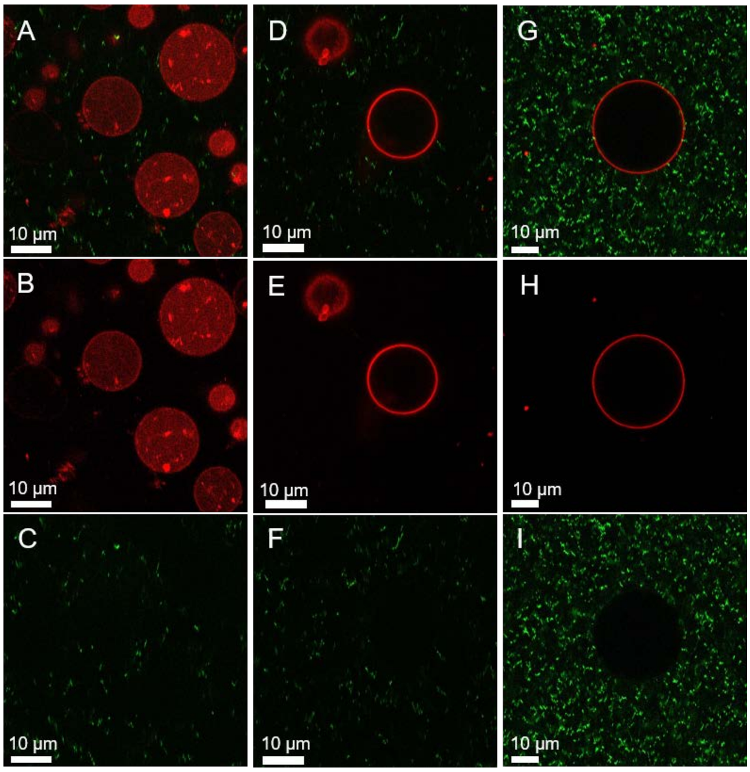

The specificity of binding of bacteriophage 9NA to its LPS receptor is illustrated for GUV preparations that contain (A-C) LPS from *S.* Typhimurium facing the GUV lumen (“inside-out” LPS-GUVs), (D-F) GUVs with LPS from E. coli HTD2158 incorporated in the outer leaflet[2] or (G-H) LPS-free POPC GUVs. A, D, G: Overlay. B, E: LPSs labelled with TexasRed C, F, I: 9NA phage labelled with SYBR Gold. H: POPC labelled with TexasRed.

**Figure S3. The amount of bound phages on GUV surface scales with LPS concentration.**

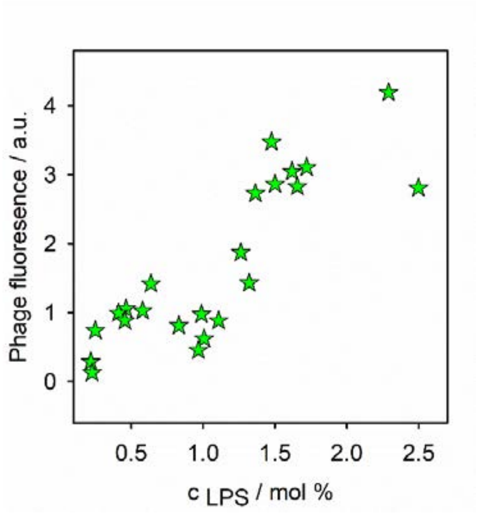

SYBR™Gold labelled *Salmonella* 9NA phage signal when localized on GUV surface plotted against LPS coverage on the GUV surface. The LPS was TexasRed-labelled and its molar concentration on GUVs quantified via a standard curve obtained with POPC-GUVs containing defined concentrations of DHPE-TexasRed.

**Figure S4. Diffusion coefficient of LPS-GUV-bound 9NA from particle tracking**

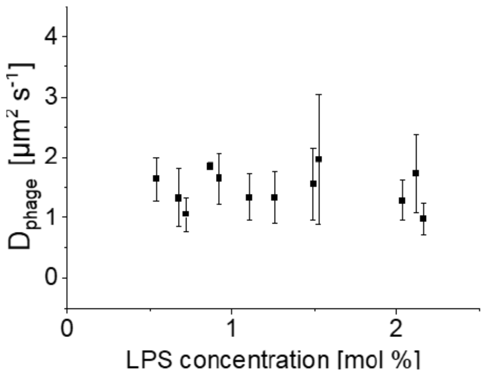

SYBR™ Gold labeled 9NA bacteriophages were co-incubated with LPS-GUVs for 15 min at room temperature. Particle tracking was conducted by recording the bacteriophage movement on top of the GUVs for 2 min with a frame rate of 25 frames per second. Mean square displacements were extracted from tracks with > 50 steps to calculate the diffusion coefficient.[3] To determine the LPS-coverage on individual GUVs, we used a calibration Texas-Red standard as previously described.[4]

### Supplementary Methods

#### Bacteriophage preparation

*Salmonella* siphovirus 9NA was propagated in *Salmonella* Typhimurium liquid cultures and purified on cesium chloride gradients as described.[1] For fluorescence labelling, 100 µl 9NA ($10^{12}$ pfu $ml^{-1}$) were mixed with 400 µl buffer (50 mM Tris-HCl, 100 mM $MgCl_2$ pH 7.6) and 2.5 µl SYBR™ Gold (10000x in DMSO) and incubated overnight at 4 °C. A PD-10 desalting column (GE Healthcare, Chicago, Illinois, USA) was equilibrated with buffer and loaded with the labeled phage sample. The sample was eluted by adding stepwise 0.5 ml bacteriophage buffer. 0.5 ml fractions were collected and spectrophotometrically checked for their phage content.

#### Preparation of LPS-GUVs from inverted emulsions

LPS of *Salmonella* Typhimurium MvP103 was prepared and fluorescently labelled with either TexasRed or AlexaFluor647 as described.[1, 4-5] An LPS mixture containing 1.2 mol % fluorescently labelled LPS was solubilized in water as 1 mg $ml^{-1}$ stock solution. LPS-GUVs were formed by an inverted emulsion technique as previously published in Stephan *et al*.[4]: First, an LPS-glucose solution was prepared. It contained 480 mM glucose, 10 mM $MgCl_2$, 10 mM Tris-HCL pH 7.6, and the 100 µg $ml^{-1}$ LPS. 250 µl of this LPS-glucose solution was placed in a 1.5 ml tube and then 200 µl lipid-free mineral oil (intermediate oil layer) was placed on top. The sample tube was mechanically agitated in a standard tube shaking rack to form an oil-in-water emulsion. After incubation at room temperature for 30 min, the sample was centrifuged for 10 minutes at 600 ×g and incubated for further 3 h or overnight. In the next step, the sample was centrifuged for 10 min at 4500 ×g to establish bulk phase separation of the oil and aqueous phase. 50 µl of 400 µM POPC lipid oil were added to fully cover the LPS-free aqueous-oil-interface. Then after 15 min incubation at room temperature, 150 µl of a 500 mM sucrose-in-oil emulsion was pipetted on top. The sample was centrifuged for 10 min at 130 ×g. The sucrose-in-oil emulsion was created by mixing of 5 µl 500 mM sucrose and 250 µl 400 µM POPC lipid oil. For phage ejection experiments, the sucrose solution contained 10 µM YO-PRO™-1 or TOTO™-3, respectively, to form GUVs with the fluorophores encapsulated in the lumen. The LPS concentration in the membrane was assessed as described, using POPC-GUVs with varying concentrations of the fluorophores as a standard.[4]

#### Purification of LPS-GUVs and delivery of bacteriophages with microfluidics

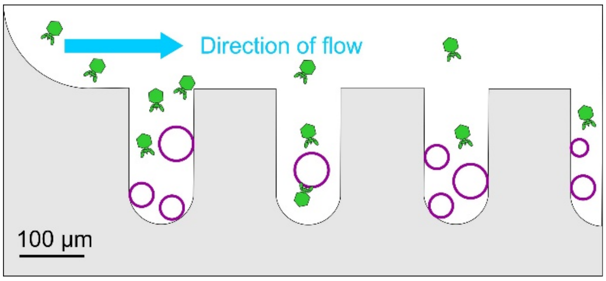

Scheme of microfluidic chip design used to expose GUVs to different solution conditions.[6] This set up was used for washing GUVs from LPS in solution or to expose them to bacteriophages. Phage particles are not drawn to scale.

The used microfluidic chip design and preparation protocol was previously described by Pramanik et al.[6] The design consists of a main channel through which the GUVs are loaded into dead-end sleeves. Once loaded into the sleeves, new solutions are flowed in the main channel and the immediate surrounding solution is exchanged by diffusion alone, which avoids exposing the vesicles to high flow rates that could cause vesicle rupture or deformation.

#### Fluorescence microscopy of bacteriophage interactions with LPS GUVs

GUVs were loaded in a microfluidic chip (see above) and washed with buffer (500 mM glucose, 4 mM $MgCl_2$, 50 mM Tris-HCl pH 7.6). 20 µl DNAse I (1 mg $ml^{-1}$) and 20 µl SYBR™ Gold labeled 9NA bacteriophage ($10^{11}$ pfu $ml^{-1}$) were mixed and loaded into the chip. Image acquisition was performed using a Leica SP8 confocal microscope with an HC PL APO CS2

63x/1.40 oil objective. The TexasRed- or AlexaFluor647 labelled lipids were excited with an OPSL laser at 552 nm or 638 nm, respectively. Emission was recorded with a HyD detector in the range of 600 to 700 nm. The SYBR™ Gold label of bacteriophages was excited with an OPSL laser at 488 nm, and the signal was recorded with a PMT detector in the range of 500 to 600 nm. The DNA bound TOTO™-3 was excited with a HeNe laser at 633 nm and the emission was detected using a PMT detector in the range of 650 to 700 nm. The YO-PRO™-1 signal from the dye bound to bacteriophage DNA was excited with an OPSL laser at 488 nm and recorded with a PMT detector in the range of 500 to 600 nm. Sequential settings were used to avoid crosstalk between channels.

### SI Movie caption

Time lapse movie (96 frames, 10 fps, 512x512 pixel) of *Salmonella* phage 9NA genome transfer into LPS-GUVs. GUVs contained *S.* Typhimurium LPS (Alexa647-labelled, magenta) on the outer surface and 10 µM of DNA intercalating dye YO-PRO™-1 (green) in the lumen (*cf.* Figure 5). The observation time was 21 min.
